# Lipidomic Fingerprints Reveal Sex-, Age-, and Disease-Dependent Differences in the TgF344-AD Transgenic Rats

**DOI:** 10.1101/2025.03.24.644980

**Authors:** Chunyuan Yin, Alida Kindt, Amy Harms, Robin Hartman, Thomas Hankemeier, Elizabeth de Lange

## Abstract

**Background:** Gathering information on Alzheimer’s disease (AD) progression in human poses significant challenges due to the lengthy timelines and ethical considerations involved. Animal AD models provide a valuable alternative for conducting mechanistic studies and testing potential therapeutic strategies. Disturbed lipid homeostasis is among the earliest neuropathological features of AD.

**Aim:** To identify longitudinal plasma lipidomic changes associated with age, sex, and AD in male and female TgF344-AD and wild-type rats.

**Methods:** A total of 751 lipids in 141 rats (n=73 TgF344-AD; n= 68 WT) were quantified at 12, 25, 50, and 85 weeks). Differential abundances of lipids were assessed using generalized logical regression models, correcting for i) age and sex, for ii) individual age groups, and iii) sex-specific differences. Predictive lipid signature models for AD were developed using stepwise feature selection for the full age range, as well as for midlife.

**Results:** Sex differences were identified among all ages in sphingomyelin (SM), phosphatidylcholine (PC), and phosphatidylethanolamine (PE) lipid classes. AD and age-related differences were found in the SM class in mid-life (25-50 weeks). Other AD and age-related differences were found in the ratios of linoleic acid and 5 of its products. Moreover, similarities in lipidomic profile changes were observed for humans and rats. The full age range and mid-life predictive lipid signatures for AD resulted in an AUC of 0.75 and 0.68, respectively.

**Conclusions:** Our findings highlight the value of lipidomic in identifying early AD-related lipid alterations, offering a promising avenue for understanding disease mechanisms and advancing biomarker discovery.

## 1. Introduction

Alzheimer’s disease (AD) is the most prevalent form of dementia in the elderly population, leading to progressive cognitive decline, functional impairment and loss of independence [1]. Over 50 million people suffered from dementia in 2020 worldwide, and this number is projected to reach 82 million in 2030 and 152 million in 2050 [2]. There is a transitional state between the cognitive changes of normal ageing and AD, known as mild cognitive impairment (MCI) [3], with a reported annual conversion rate from MCI to AD of 10-15%, compared with cognitive normal subjects at a rate of 1-2% per year [4, 5]. Lifestyle changes and treatment could significantly delay the onset of AD once MCI is diagnosed [6], thus stressing the importance of early identification of MCI. The U.S. Food and Drug Administration (FDA) approved Aduhelm (aducanumab), targeting amyloid-beta plaques, in June 2021 and Leqembi® (lecanemab, Eisai/Biogen), an anti-amyloid-beta protofibril antibody, in July 2023 for the treatment of early AD [7, 8]. While these are the first approved AD drug treatments proven to slow the progression of the disease, there is still substantial potential for improvement in the treatment of AD, especially at the earlier stage of AD. To that end, there is a critical need for information and a deeper understanding of the processes associated with the onset and initial phases of AD. The challenge lies in acquiring more detailed mechanistic information on the time course and interrelationships of the processes that drive the onset and early development of MCI and AD.

Cerebrospinal fluid (CSF) testing and non-invasive brain scans using either positron emission tomography (PET) or structural magnetic resonance (MRI) are considered two core approaches for the reflection of different of AD pathology. Several CSF biomarkers for AD, including amyloid-β42 (Aβ42), Aβ42/40, phospho-tau181 (P-tau181), neurogranin concentration, and neurofilament light (NfL) concentration have been found [9-13]. However, CSF testing is invasive and sometimes painful, and brain scans are too expensive for routine care, in which blood sampling is cheaper and more readily accessible, especially for elderly.

Metabolomics enables the systematic examination of unique metabolomic signatures arising from the body’s functioning under various conditions, including both healthy and diseased states. These signatures can be regarded as biomarkers indicative of normal biological processes, pathological changes, or a body’s response to therapeutic interventions. Lipids are essential nutrients that play key roles in biological and physiological processes, such as cell signaling and impulse conduction [14]. Lipidomics has been widely used to explore the relationship between lipids and AD progression [15-18]. A recent review summarized the current state of metabolomics research on AD, focusing on studies conducted in human and animal models using plasma, CSF, and brain samples [19]. Monitoring blood or plasma levels for potential biomarkers is both feasible and ongoing in multiple medical centers [20-24]. However, longitudinal studies that integrate data from multiple body sites (brain, CSF, and blood) remain scarce. These studies are crucial for understanding the interrelationships among various AD processes [25]. Such information can be difficult to obtain in human studies due to the long observation time involved, as well as the inability to directly sample human brain.

Since gathering such data in humans presents significant challenges, a humanized animal model of AD, such as the TgF-344 AD rats, could be particularly useful as they provide fresh perspectives on disease biomarker patterns and biological pathways. Such insights not only support AD stage diagnosis in humans but also pave the way for discovering future therapeutic targets for AD and evaluate treatment options. A thorough characterization of the lipid pathways involved in AD progression in this animal model has to date not been performed. The shorter life span of these rats allows for a much shorter investigative period which eases the identification of processes associated with the initiation and development of AD. The TgF344-AD transgenic rat model overexpresses the human amyloid precursor protein (APP) with the Swedish mutation and PSEN1 with the Δ exon 9 mutation that results in an age-dependent AD pathology, including the accumulation of amyloid plaques, hyperphosphorylated tau, neuronal loss, gliosis, neuroinflammation, and progressive memory impairment. Previous literature has shown that memory impairment of TgF344-AD rats starts around 24 weeks of age, with significant cognitive deficits observed around 60 weeks compared to WT rats [26-30]. These features make the TgF344-AD rat model an attractive preclinical AD model for elucidating early changes in metabolism associated with the disease. In our laboratory, a biobank was created with body fluids and tissues, in a longitudinal manner, from male and female AD rats (male = 33, female = 40) and their wild type littermates (male = 35, female = 33), to investigate their compositions in an age, sex and genetic background dependent manner.

In this study, we used UPLC-MS to relatively quantify 751 lipids, covering 13 classes, in 73 plasma samples of TgF344-AD and 68 WT rats obtained at 12, 25, 50 and 85 weeks. The aim was to identify associations of lipid changes associated with normal ageing, sex, and AD. A lipid fingerprint was developed using stepwise feature selection to distinguish AD from WT using age and sex as covariates.

## 2. Materials and methods

### 2.1. Animals and plasma collection

Heterozygous TgF344-AD rats expressing mutant human APP_sw_ and PS1ΔE9 genes and their age-matched wild-type Fischer344 littermates were used in this study. This strain was co-injected with two transgenes which integrate at the same chromosomal site: the first transgene contains human amyloid beta precursor protein with the “Swedish” mutation. The second transgene contains human PSEN1 with a deletion of exon 9 (PS1ΔE9). Both transgenes are driven by the mouse prion promoter [26]. The rats were purchased from the Rat Resource & Research Center of the University of Missouri (Columbia, MO) and bred in the Animal Research Facility of Leiden University. Animal protocols were approved by the Leiden University Animal Welfare Body Leiden (AWB: AVD1060020171766).

A longitudinal animal study was performed with a total of 141 male and female TgF344-AD rats and wildtype littermates (WT) over four ages (12, 25, 50, and 85 weeks). The overall experimental design and group information is show in Figure-1. All rats were bred in a facility of Leiden University in a light and dark 12h/12h alternate cycle with regulated temperature (21±2°C) and humidity (40-60%) and fed the same rodent maintenance diet throughout their stay. Blood samples were drawn directly from the left ventricle of the heart and transferred to ethylenediaminetetraacetic acid (EDTA) tubes containing the anticoagulant EDTA. These samples were then centrifuged at 2300G for 10 minutes at 4°C, after which the supernatant (plasma) was transferred and aliquoted into new tubes for storage at -80°C. Additionally, various tissues were collected, weighed, and stored at -80°C for further analysis.

**Figure 1.**
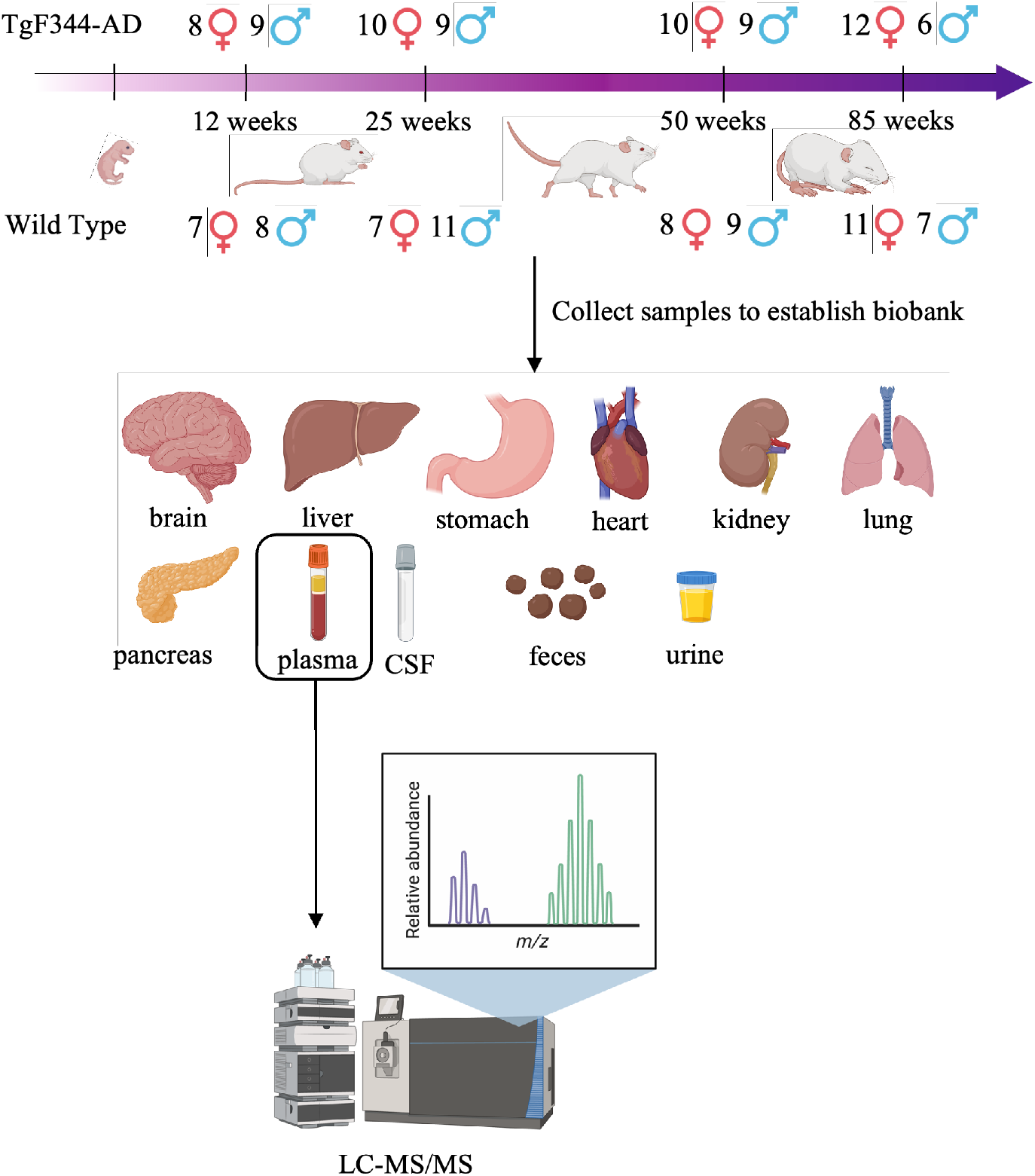
The overall experimental design and group information. Schematic representation of the experimental design, including the time points (12, 25, 50, and 85 weeks) for sample collection from TgF344-AD and WT rats. Sample collection include plasma, CSF, feces, urine, and tissues (brain, liver, stomach, heart, kidney, lung, and pancreas) to establish a biobank for further analysis. Plasma lipid profiles were analyzed using LC-MS/MS.

### 2.2. Plasma lipid analysis

#### 2.2.1. Lipid platform

EDTA Plasma samples were prepared as previously reported [31]. Briefly, plasma samples were prepared by liquid-liquid extraction using MTBE:MeOH:Water (10:3:2.5, v/v/v), and analyzed by HILIC UHPLC-MS/MS using a Luna amino column (100 mm × 2 mm, 3 μm, Phenomenex). Mass spectrometry was conducted using a Sciex QTRAP 6500 MS (Sciex, Framingham, MA, USA). ESI-MS was performed with polarity switching and multiple-reaction-monitoring (MRM). Targeted lipid classes included ceramides (Cer), cholesterol esters (CE), diglycerides (DG), lysophosphatidylcholine (LPC), lysophosphatidyl-ethanolamine (LPE), phosphatidylcholine (PC), phosphatidylethanolamine (PE), sphingomyelin (SM), triglycerides (TG), hexosylceramides (HexCer), and lactosylceramides (LacCer). These lipid classes were part of the standard mix from the system suitability kit for the Lipidyzer platform (part no. 5040407) purchased from AB Sciex (Framingham, MA, USA). The acquired LC-MS data were processed using Sciex MultiQuant software (v3.0.2, Sciex), integrating the assigned MRM peaks and further correcting according to the peak areas of matched internal standards. In-house quality-control software was utilized to assess and correct the analytical performance based on study QC replicates, blank samples, and internal standards. A total of 628 lipid features passed quality control (RSD QC <30%) and were utilized in the statistical analysis.

#### 2.2.2. Signaling lipid platform

EDTA Plasma samples were prepared as previously described using liquid-liquid extraction (LLE) with butanol and methyl tert-butyl ether (MTBE) [32]. In short, the organic phase was dried and reconstituted for injection into two separate RPLC-MS/MS methods (high and low pH), enabling the analysis of 260 metabolites, including oxylipins (isoprostanes, prostaglandins, and other oxidized lipids), free fatty acids, lysophospholipids, sphingoid bases (C16, C18), platelet activating factors (C16, C18), endocannabinoids, and bile acids. High pH analysis was performed using a Shimadzu LCMS-8050 triple quadrupole mass spectrometer with a Kinetex EVO column, while low pH analysis was conducted using a SCIEX QTRAP system with an Acquity UPLC BEH C18 column. Both methods utilized polarity switching and dynamic MRM mode for analyte monitoring. Data were processed using LabSolutions Insight (Version 3.3, Shimadzu) and SCIEX OS software (version 2.1.6.59781). In-house quality-control software (mzQuality) was utilized to assess and correct the analytical performance based on study QC replicates, blank samples, and internal standards. A total of 123 lipid features with RSD QC < 30% measured by this method passed quality control and were utilized in the statistical analysis.

### 2.3. Statistics analysis

All statistical tests were performed using R studio software (Version 4.2.1). The lipids that passed quality control were log_2_ transformed prior to the analysis. Scaling was performed to calculate the mean aggregated z-score for class means, and for visualization where appropriate. Biologically relevant ratios were calculated by dividing products by their precursors for enzyme-dependent conversions, resulting in a comprehensive dataset of 825 lipids species (628 from the lipid platform and 123 from the signaling lipid platform) and ratios (n=74). For a detailed list of the calculated ratios, see Supplementary Table 1.

**Table 1.** The estimated coefficients and p value of predictors in the full age range model-After Intercept, Age and Sex, the outcomes, are displayed in order of increasing p value.

|  | Estimate | p value |
| --- | --- | --- |
| (Intercept) | 3.92 | 1.13E-03 |
| Age | -0.05 | 9.94E-03 |
| Sex (relative to female) | -2.87 | 1.54E-02 |
| PC (16:0/19:0) | -2.19 | 2.00E-04 |
| PE (18:1/20:3) | 1.93 | 7.29E-04 |
| SM (d29:1) | 1.74 | 2.09E-03 |
| PE (O-18:0/20:2) | -1.62 | 4.20E-03 |
| LPI (18:0) | 1.80 | 4.32E-03 |
| CE (18:1) | -1.40 | 4.74E-03 |
| GCDCA | -1.53 | 8.32E-03 |
| PE (20:4/20:4) | -1.53 | 9.01E-03 |
| 12-S-HEPE/12-HETE | 1.15 | 1.84E-02 |
| LPC (20:5) | 1.27 | 2.01E-02 |
| 9,12,13-TriHOME/LA | -1.30 | 2.13E-02 |
| GDCA | 1.24 | 2.17E-02 |
| cLPA (18:0) | 1.09 | 2.48E-02 |
| S-1-P (18:2) | 1.21 | 3.26E-02 |
| TG56:2-FA20:1 | -1.06 | 4.16E-02 |
| TG58:10-FA20:5 | -0.85 | 6.05E-02 |
| PE (O-18:0/18:0) | 0.86 | 6.36E-02 |
| LPA (18:0) | -1.04 | 6.47E-02 |
| 15-HETE | 1.12 | 6.82E-02 |
| LPE (20:5) | -1.01 | 7.30E-02 |
| PC (O-38:6/18:2) | 0.82 | 8.21E-02 |
| PE (18:2/18:2) | 0.98 | 8.21E-02 |
| SM (d30:0) | -0.84 | 8.82E-02 |
| 15-HETrE/DGLA | -1.04 | 9.17E-02 |
| TDCA/DCA | 0.71 | 1.08E-01 |
| DHA | -0.87 | 1.12E-01 |
| S-1-P (18:0) | -0.84 | 1.32E-01 |
| CE (22:5) | -0.66 | 1.44E-01 |
| LPG (16:1) | 0.67 | 1.46E-01 |
| AEA | -0.57 | 1.47E-01 |
| LPS (18:1) | -0.80 | 1.52E-01 |

#### Univariate analysis

Differential lipid abundances in TgF344-AD rats compared to WT rats were analyzed using generalized logistic regression models (GLM). Included confounders were: (i) age and sex in the overall analysis, (ii) individual age groups in sex-specific models, and (iii) sex within different age models. To account for multiple comparisons, p-values were adjusted using the Benjamini-Hochberg method.

#### Predictive modeling

A full age range and a midlife predictive model for AD were developed using a combination of identifying highly correlated variables and a stepwise feature selection method. We first applied the “FindCorrelation” function, a pairwise correlation method, to identify and remove highly correlated lipids (r > 0.65) to avoid collinearity in the full age range model; for the midlife model, a lower threshold (r > 0.50) was applied. Based on these variables, we then employed a stepwise feature selection model to identify the lipids most strongly associated with AD. The model was refined by selecting features that minimized the Akaike Information Criterion (AIC) scores, an estimator of in-sample prediction error that is comparable to adjusted R-squared measures in common regression methods [33]. The feature selection algorithm applied performed an addition or removal of variables at each step. The model resulting from this algorithm was refined by iteratively replacing the previously removed strongly lipids to derive the final model. For reproducibility, a seed of 99 was set in stepwise feature selection method.

The R packages dplyr (version 1.1.2), reshape (version 0.8.9), stringr (version 1.5.0), tidyverse (version 1.3.2), caret (version 6.0-94) was used. For visualizations R packages pheatmap (version 1.0.12), ggrepel (version 0.9.2), ggplot2 (version 3.4.2), ggpubr (version 0.5.0), geomtextpath (version 0.1.1), plotrix (version 3.8-2), ggrain (version 0.0.3) were used.

## 3. Results

### 3.1. Principal component analysis of the plasma lipidomic data

A principal component analysis (PCA, Figure 2A) demonstrated an age-related separation along the second principal component (PC2 10.01% explained variance), where both the WT and TgF344-AD groups exhibit a positive shift with increasing age. The PCA loading plot (Figure 2B) reveals that lipid classes such as eicosanoids, endocannabinoids, and SMs show a positive association with PC2, while lysophospholipids and eicosanoids exhibit a negative association. In contract, PC1 (50.32% variance explained) primarily captures variance related to lipid classes such as TGs, PCs and PEs, which are distributed along the positive direction of the PC1 axis. The PCA loading plot further highlights clusters of PCs, SMs, and TGs, suggesting potential similarities in their roles or metabolic pathways. Lipids located far from the origin, particularly TGs and PCs in PC1 axis, emerge as prominent contributors to the variance and may serve as key factors distinguishing TgF344-AD from WT groups. This distinction needs further investigation to clarify the roles of lipid classes in driving AD-specific metabolic changes versus general age-related shifts.

**Figure 2.**
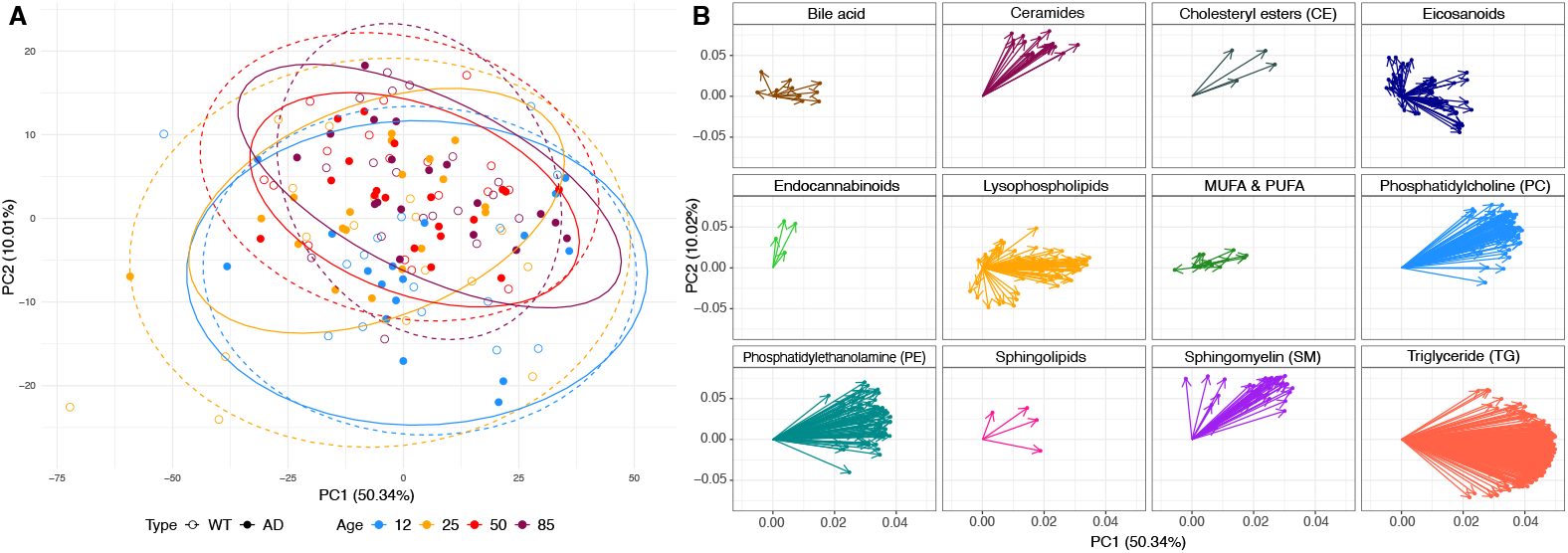
Principal component analysis (PCA) plot of 141 rat plasma samples. [A] Scores plot showing samples from 12 weeks (blue markers), 25 weeks (orange markers), 50 weeks (red markers) and 85 weeks (dark red markers), based on all measured lipids (log_2_ transformed and auto scaled). Data points from wild-type (WT) group are depicted in open circle shape and are enclosed within dotted lines, while data points from transgenic TgF344-AD rats (AD) are depicted in closed circles shapes and are enclosed within solid lines. [B] PCA loadings plot visualizes how lipids contribute to the principal components (PC1 and PC2). The distance of these points from the origin indicates how much the lipid contributes to the variance captured by the principal components. Each arrow represents a lipid’s loading showing the direction and strength of the lipids’ influence on the principal components. The longer the arrow, the greater the influence of that lipid on the respective principal component.

### 3.2. Differentially abundant lipids between TgF344-AD and WT rats

To identify differences in the lipidome of TgF344-AD and WT rats, a generalized logistic model was calculated correcting for age and sex of the rats (Figure 3A). Here, 40 of 825 lipids, where 25 TGs, SM(d36:4), SM(d34:3), SM(d36:3), SM(d34:2), SM(d33:2), SM(d36:2), SM(d44:6), PC(16:1/18:0), PC(16:0/22:5), PE(O-16:0/22:5), PE(P-16:0/22:5), 15-HEPE, 15-HEPE/15-HETE, LEA and alpha-LA, were increased in TgF344-AD rats, while only glycocholic acid (GCA) was decreased. The most increased lipid was SM(d36:4) (*p*=2.5E-03, estimate=0.26), followed by alpha-LA (*p*=6.8E-03, estimate=0.42) and PE(O-16:0/22:5) (*p*=7.3E-03, estimate=0.25).

**Figure 3.**
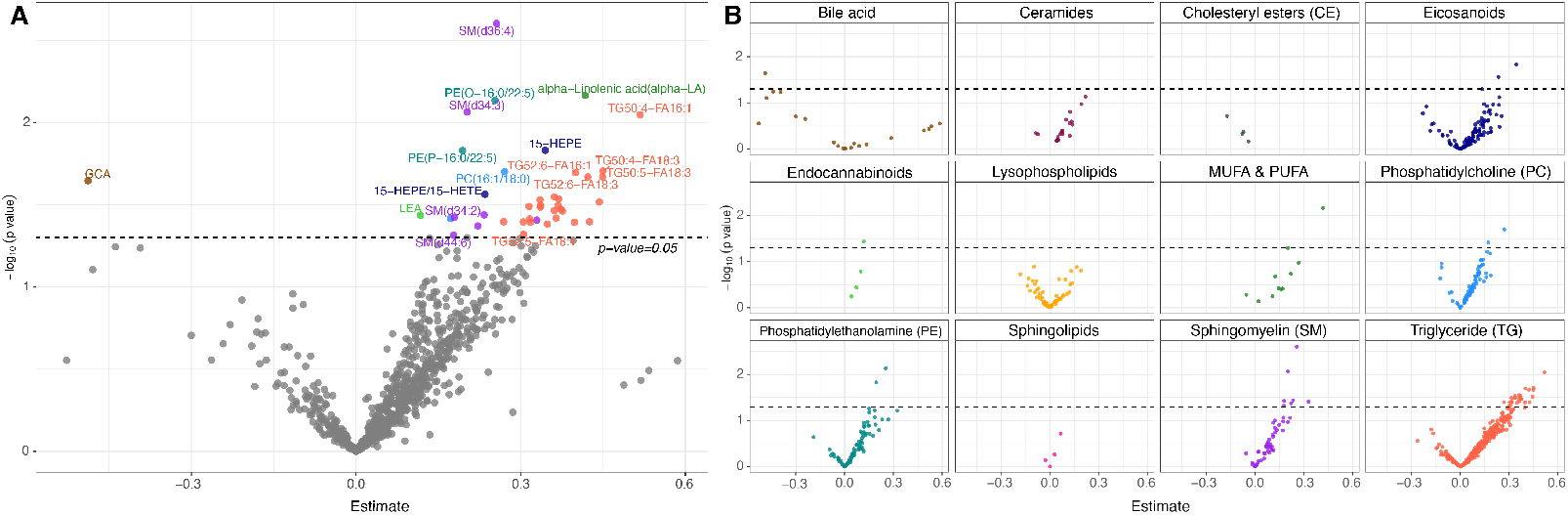
[A] Volcano plot of the detected lipids showing GLM results of all TgF344-AD rats versus all WT rats (x-axis: estimate; y-axis: -log10(p value)). Lipids with p-value <0.05 are marked with different colors representing the class to which individual lipid belong. The dotted line represents the significance threshold (*p* value = 0.05). [B] Lipid class-specific volcano plots showing the distribution of individual lipid species across different classes, including bile acids, ceramides, cholesteryl esters (CE), eicosanoids, endocannabinoids, lysophospholipids, monounsaturated fatty acids and polyunsaturated fatty acids (MUFA&PUFA), phosphatidylcholines (PC), phosphatidylethanolamine (PE), sphingolipids, sphingomyelins (SM), and triglycerides among others. Each panel illustrates the significance (−log10(p value)) and effect size (estimate) for lipids within the respective class.

To further explore class-specific lipid changes, Figure 3B presents volcano plots stratified by lipid classes, including bile acids, ceramides, CE, eicosanoids, endocannabinoids, lysophospholipids, monounsaturated fatty acids and polyunsaturated fatty acids (MUFA & PUFA), PC, PE, sphingolipids, SM, and TG. Each panel illustrates the significance (−log10(p value)) and effect size (estimate) for individual lipids within their respective classes. Notably, TG and SM classes showed the largest number of altered lipids, consistent with the overall findings in Figure 3A.

### 3.3. Lipid abundances change earlier in TgF344-AD rats than in WT rats

As reported in a previous study, the TgF344-AD rat model shows a trend toward memory impairment from 24 weeks of age onwards, with significant cognitive deficits observed starting around 60 weeks compared to WT rats [26]. Therefore, we focused our investigation on the 25-week and 50-week (Figure 4) stages. At 25 weeks (Figure 4A), sphingolipids showed elevated levels in the TgF344-AD group, with several sphingolipid species showing upregulation, e.g., SM(d36:2), *p* = 4.45E-03; SM(d36:4), *p* = 3.97E-03. By 50 weeks (Figure 4B), there is a shift, where additional sphingolipids are more prominent and still high in TgF344-AD, e.g., SM(d40:7), *p* = 1.87E-02 and SM(d37:2), *p* = 8.08E-03. Notably, DHA exhibited changes at both time points, appearing on the negative side of the x-axis at 25 weeks and on the positive side at 50 weeks.

**Figure 4.**
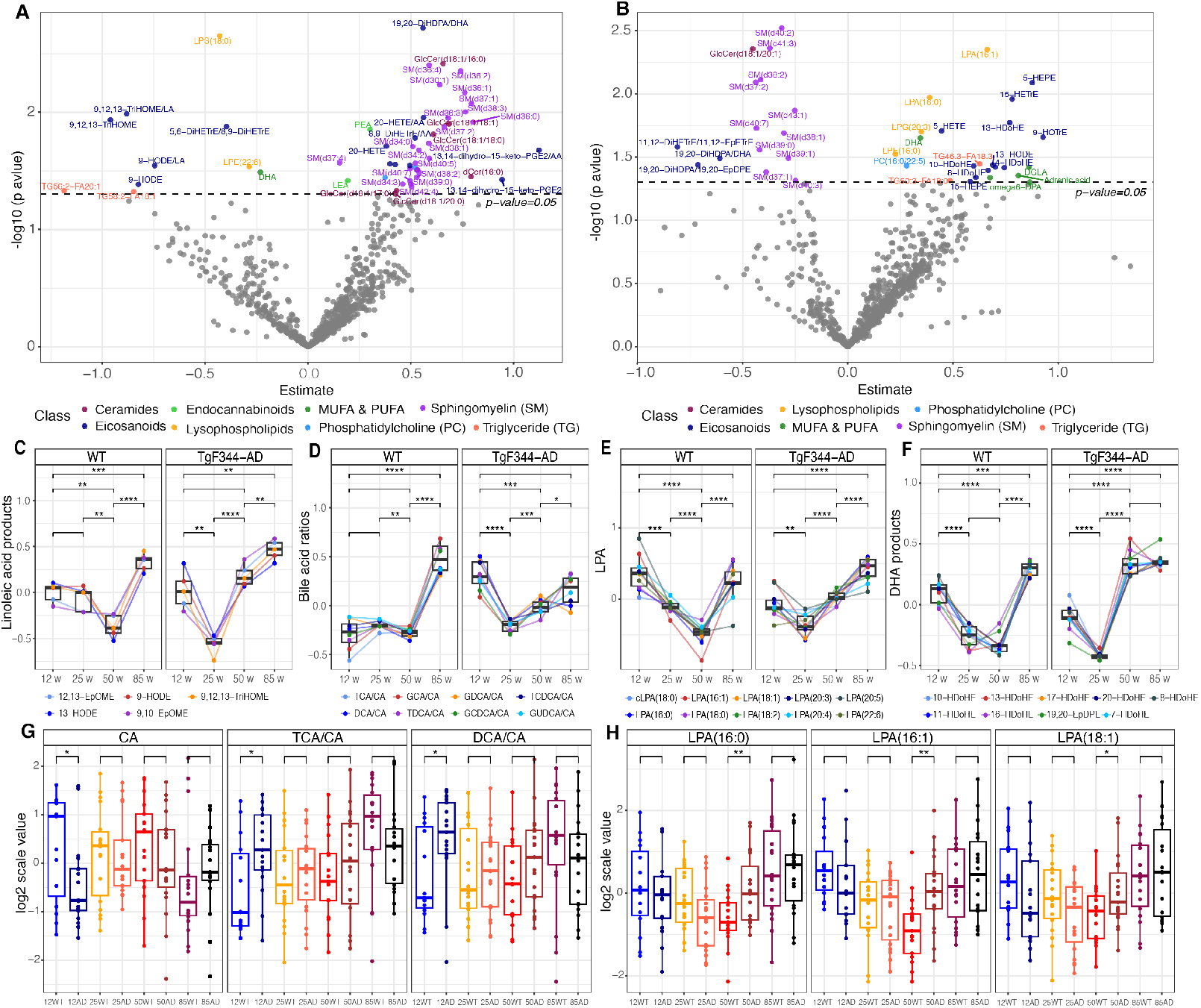
Comparative analysis of metabolic lipid levels in TgF344-AD rats and WT rats. [A] Volcano plot of generalized logistic model (GLM) results for 25-week-old TgF344-AD rats versus WT rats. [B] Volcano plot of GLM results for 50-week-old TgF344-AD rats versus WT rats. [C-F] Trends in key lipid classes and ratios, including linoleic acid metabolites, bile acids ratios, lysophosphatidic acids (LPAs), and DHA-derived metabolites, across different age points. [G-H] Boxplots of specific lipids (e.g., CA, DCA/CA, TCA/CA, and LPAs), illustrating differences between TgF344-AD and WT rats across different age points.

Furthermore, these lipid alterations extend across multiple age points and lipid classes, as shown in Figures 4C-4H. In Figure 4C, five linoleic acid metabolites (12,13-EpOME, 9-HODE, 13-HODE, 9,10-EpOME, and 9,12,13-TriHOME) in both WT and TgF-344 AD rats exhibited a similar trend where levels decreased around 25-50 weeks but rose again by 85 weeks. However, the TgF-344 AD rats experienced more notable changes, suggesting that metabolic activity is changing faster in the TgF344-AD rats when comparing to the WT rats. A similar pattern is observed in the ratios of secondary and primary bile acids with respect to cholic acid (CA), as shown in Figure 4D. Across the different age groups, a gradual reduction in levels is observed until 50 weeks in WT rats and 25 weeks in TgF-344 AD rats, followed by a sharp increase. The TgF-344 AD group shows these changes earlier and more drastically, indicating accelerated bile acid metabolism in TgF-344 AD compared to normal ageing. The LPA panel (Figure 4E) shows changes in lysophosphatidic acids (LPAs) of varying carbon chain lengths. In both WT and TgF-344 AD groups, before increasing at 85 weeks, the WT group dropped at 25 weeks and AD group dropped at 50 weeks. Additionally, in the TgF-344 AD rats, the increase is more pronounced, again suggesting that AD accelerates LPA metabolism when comparing to normal aging. The DHA panel (Figure 4F) presents the fluctuations of nine DHA-derived metabolites. Both groups show a dip around mid-life (25-50 weeks) and a rebound by 85 weeks. Additionally, two ratios of secondary BAs to CA (DCA:CA and TCA:CA) were higher in the AD group at 12 weeks (Figure 4G). Three LPAs (C16:0, C16:1, C18:1) showed higher levels in the AD group at 50 weeks (Figure 4H).

### 3.4. Sex associated differences in lipids

Next, we tested whether sex-associated differences in plasma lipid levels differ between TgF344-AD and WT rats (Figure 5A). From the complete dataset (n=141), 54 out of 825 lipids were found to be associated with sex after adjusting for age (Supplementary Table-2). These included 4 bile acids, 6 SMs, 18 TGs, 6 PCs, 3 PC-O species and 2 eicosanoids, all of which were more abundant in male rats (highlighted in blue). Meanwhile, 6 TGs along with alpha-linoleic acid, oleic acid, PE(O-16:0/22:5), PC(16:1/18:0), PC(O-38:6)-FA18:1, 15-HEPE/15-HETE, and 12S-HEPE/12-HETE were more abundant in female rats (highlighted in yellow). Both females and males exhibit associations with TGs, with the female group exhibiting more long-chain TGs, particularly those with a total of 60 carbons in their acyl chains (e.g., TG-60), while the male group exhibits more short-chain TGs, especially those with 50 carbons in their acyl chains (e.g., TG-50).

**Figure 5.**
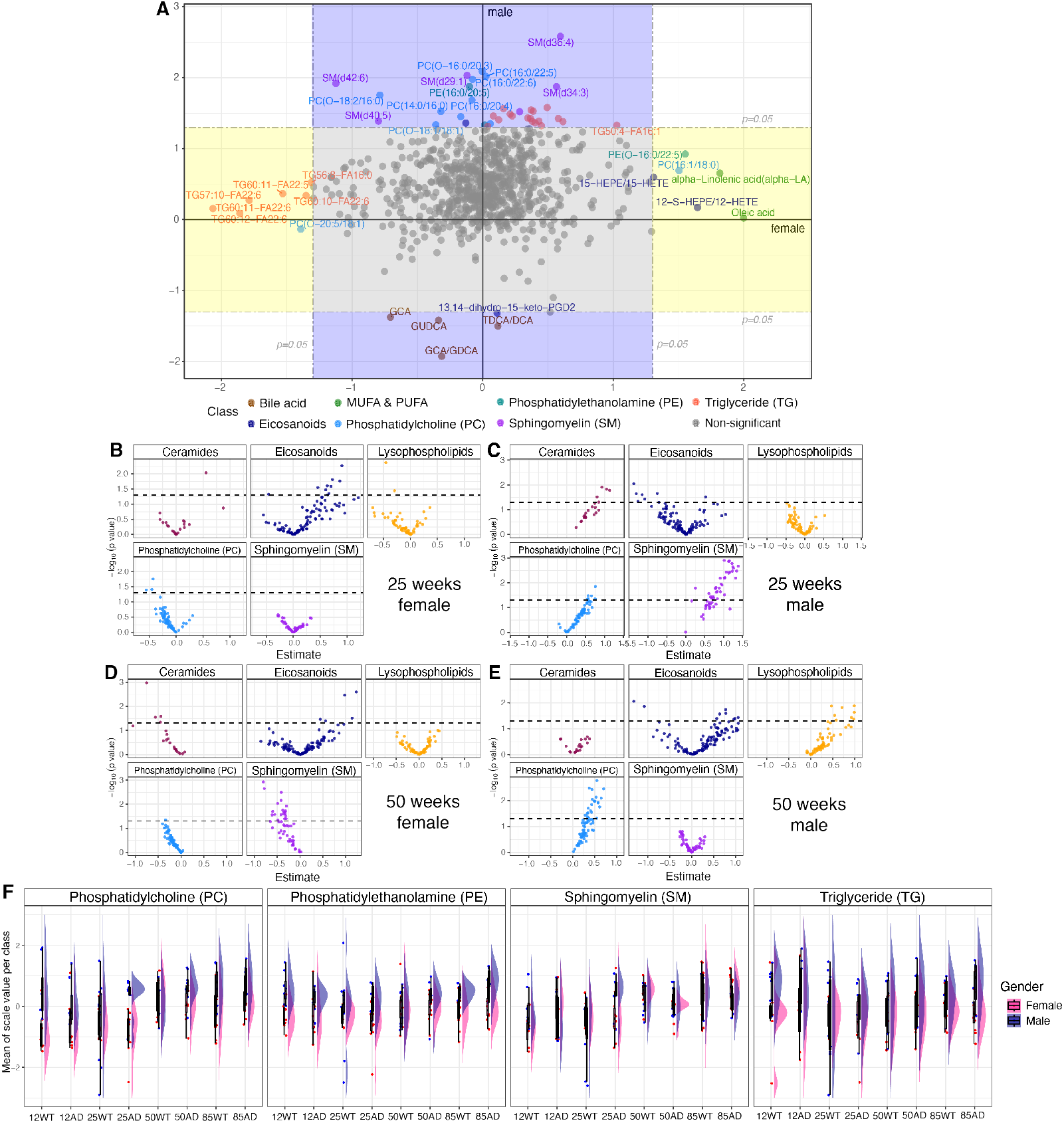
[A] Directed p-value plot summarizing generalized logistic model (GLM) results for female (x-axis) verses male (y-axis). Points in the blue region represent AD-specific lipids with changes in males, while points in the yellow region represent AD-specific lipids with changes in females. Gray points indicate *p* > 0.05. A directed p-value is the -log_10_ p-value times the sign of the estimate. [B-E] Sex-based lipid divergence in [B] 25 weeks female rats, [C] 25 weeks male rats, [D] 50 weeks female rats; and [E] 50 weeks male rats. [F] Raincloud plot shows lipid class distribution differences between male and female across ages and disease groups. The x-axis represents different time points and groups, while the y-axis indicates the scaled and averaged relative ratios per lipid class (PC, PE, SM, and TG).

This sex-based divergence in lipid profiles is further explored at 25 and 50 weeks in TgF344-AD rats compared to WT rats. In Figure 5B (25-week female) and Figure 5C (25-week male), notable differences are observed in ceramides, eicosanoids, PCs, and sphingomyelin, with sphingomyelin being elevated (*p*<0.05) in the AD male group. By 50 weeks, the metabolic differences became more pronounced (Figure 5D&E). In Figure 5D (50-week female), eicosanoids continued to show alterations, and sphingomyelin in females showed a notable increase. In contrast, Figure 5E (50-week male) highlights even stronger differences in phosphatidylcholines. Taken together, these findings highlight sex-dependent differences in lipid metabolism between TgF344-AD and WT rats, particularly in sphingomyelin, phosphatidylcholines, and eicosanoids.

A raincloud plot (Figure 5F) reveals distinct patterns in lipid class distribution between male and female groups across different age and disease groups. Especially, the PC class shows the most notable sex differences, particularly in the 12 weeks old WT rats and 25 weeks old TgF344-AD rats, where male and female results barely overlap. In these groups, the median and distribution of males and females are clearly separated, indicating potential lipid metabolism differences associated with sex at these two time points. Similarly, SMs display notable sex differences among 25 weeks old TgF344-AD rats.

Across the other age groups and lipid classes, such as PE and TG, the results for males and females show more alignment; however, the sex-based variations in the distributions can still be observed. These differences, though less significant, may still point to sex-based variability in lipid metabolism across ageing and disease progression.

### 3.5. Development of a predictive model for AD by stepwise regression

The full age-range model, which includes lipid data across all ages (12, 25, 50, and 85 weeks) along with age and sex factors, distinguishes between TgF344-AD and WT rats with an AUC of 0.75, a sensitivity of 0.71, a specificity of 0.66, an accuracy of 0.69, a PPV of 0.69, and F1-score of 0.70 (Figure 6A), which is better than the lipid only model (AUC = 0.71, sensitivity = 0.67, specificity = 0.62, accuracy = 0.65, PPV = 0.65, F1-score = 0.66). The leave-one-out validation performed using 141 rats (AD = 73; WT = 68) further supports the robustness of this multivariate approach. Estimates of the variables and their statistical significance (p-values) in the final model are shown in Figure 6B, where different colors represent different lipid classes. The size of each dot indicates the p-value, with larger dots representing more statistically significant predictors. Age, sex, PC (16:0/19:0), PE (18:1/20:3), SM (d29:1), PE-O (18:0/20:2), LPI (18:0), CE (18:1), GCDCA, and PE (20:4/20:4) show strong contributions to the model, as seen from their larger effect sizes and smaller p-values (Table 1).

**Figure 6.**
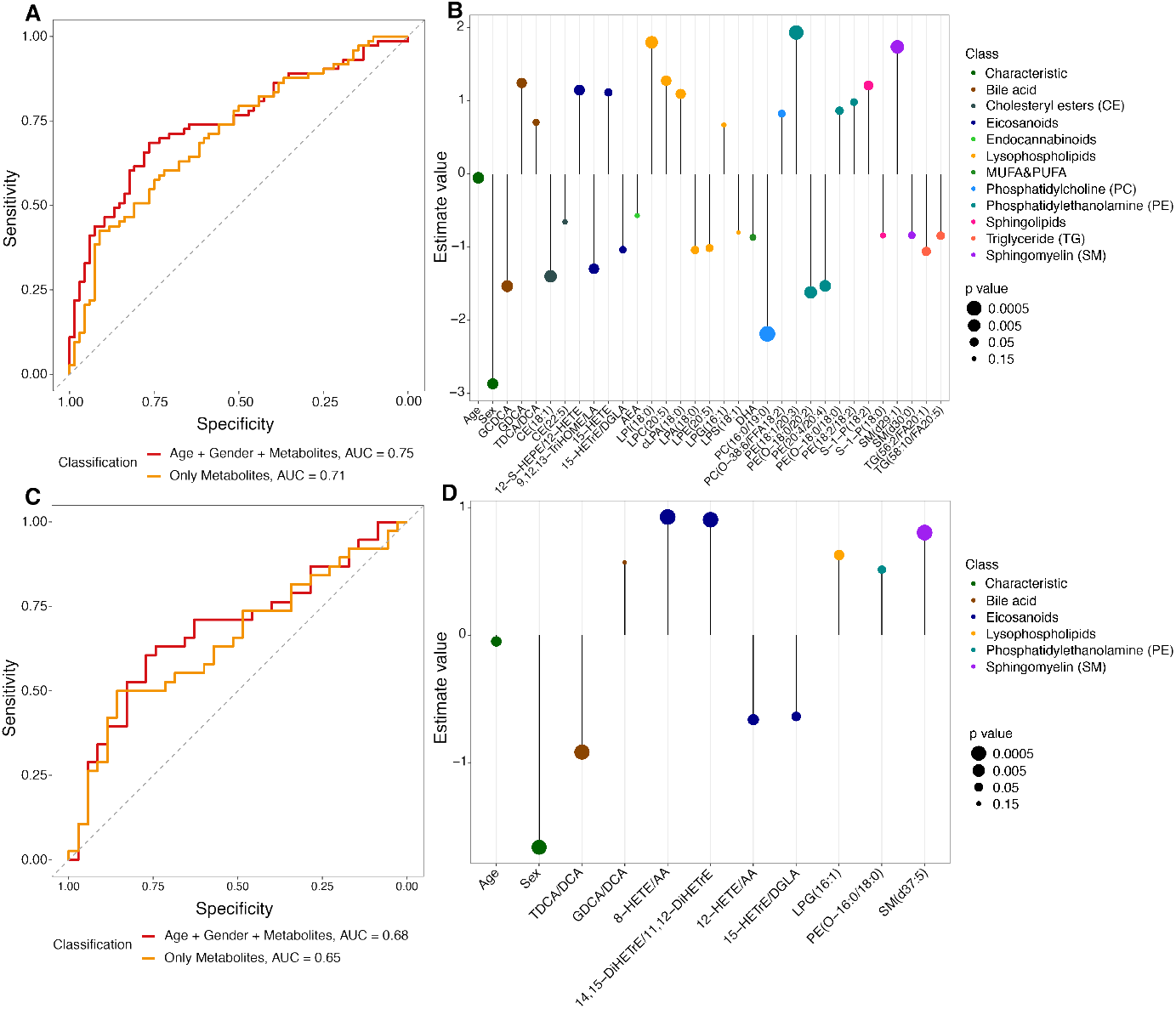
[A] Multivariate model discriminates between TgF344-AD and WT rats. Leave-one-out validation on 141 rats (AD=73; WT=68) shows improved performance with “age + sex + lipids” (red, AUC = 0.75) compared to lipids only (orange, AUC = 0.71). [B] Coefficients and p-value of predictors in the full age-range model, grouped by lipid classes and characteristics, with larger dots indicating higher significance. [C) Midlife model (25-week and 50-week) validation on 73 rats (AD=38; WT=35) shows “age + sex + lipids” (red, AUC = 0.68) outperforming lipids only (orange, AUC= 0.65). [D] Forest plot of coefficients and p-value in the midlife model.

We also developed a midlife lipid signature model for the 25 and 50-week time points. This model distinguishes between AD and WT rats with an AUC of 0.65, sensitivity of 0.63, specificity of 0.54, accuracy of 0.59, PPV of 0.6, and an F1-score = 0.62. This model was further improved by adding age and sex as covariates to have an AUC of 0.68, a sensitivity of 0.63, a specificity of 0.69, an accuracy of 0.66, a PPV of 0.69, and F1-score of 0.66 (Figure 6C). The estimates and the significance of predictors in the 25-week and 50-week model are shown in Figure 6D and Table 2. Here, key lipid classes such as bile acids, eicosanoids, lysophospholipids, sphingolipids, and PE still contribute to the models.

**Table 2.**
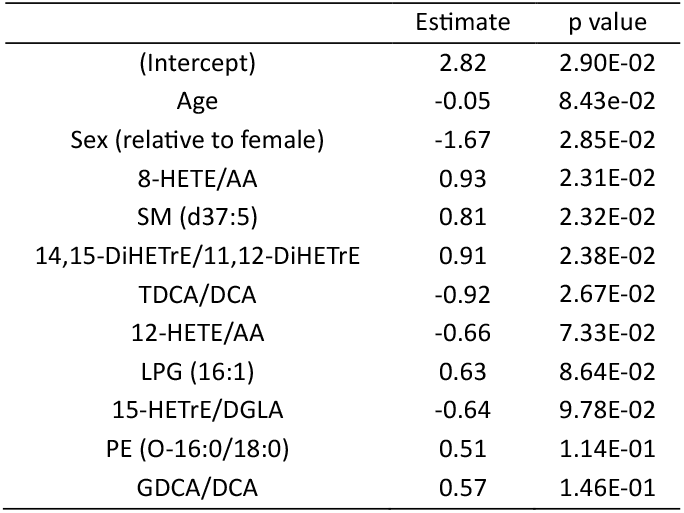
The estimated coefficients and p value of predictors in the midlife lipid signature model. After Intercept, Age and Sex, the outcomes, are displayed in order of increasing p value.

|  | Estimate | p value |
| --- | --- | --- |
| (Intercept) | 2.82 | 2.90E-02 |
| Age | -0.05 | 8.43e-02 |
| Sex (relative to female) | -1.67 | 2.85E-02 |
| 8-HETE/AA | 0.93 | 2.31E-02 |
| SM (d37:5) | 0.81 | 2.32E-02 |
| 14,15-DiHETrE/11,12-DiHETrE | 0.91 | 2.38E-02 |
| TDCA/DCA | -0.92 | 2.67E-02 |
| 12-HETE/AA | -0.66 | 7.33E-02 |
| LPG (16:1) | 0.63 | 8.64E-02 |
| 15-HETrE/DGLA | -0.64 | 9.78E-02 |
| PE (O-16:0/18:0) | 0.51 | 1.14E-01 |
| GDCA/DCA | 0.57 | 1.46E-01 |

## 4. Discussion

In this study, we examined the plasma lipidome of male and female TgF344-AD and wild-type rats at 12, 25, 50, and 85 weeks. Predictive models for AD were developed by stepwise feature selection to distinguish AD from WT using age and sex as covariates using the full age range as well as focusing on the midlife stage of rats. Further, AD, sex and age-associated differences were found for several lipid classes. The research presented here provides fundamental information on lipid dysregulation during various stages of TgF344-AD rats and normal ageing rats. This study also highlights potentially shared mechanistic pathways between humans and the AD rat model, supporting its translational relevance.

Our research focused on lipids. Lipids have been increasingly recognized for their significant role in AD, particularly in relation to neurodegeneration and inflammation [34]. However, so far, the time course of lipid dysfunction and AD progression remains unclear. By studying these profiles in a longitudinal fashion in rats, we were able to identify distinct lipidomic fingerprints associated with the progression of AD. Multiple lipids were found to be changed when comparing the AD with WT rats. In the AD rats, the most increased lipid was SM(d36:4), followed by alpha-LA, and then followed by PE(O-16:0/22:5). Three LPAs (C16:0, C16:1, C18:1) showed higher levels (p-value < 0.05) in the AD group at 50 weeks when compared to WT. Interestingly, this result is consistent with the findings from human AD study [35], which reported that LPAs (C16:0, C16:1) were significantly linked to AD biomarkers in CSF (Aβ-42, p-tau, and total tau), suggesting that LPAs play a role in AD. Additionally, the ratios of secondary to primary BAs (DCA:CA and TCA:CA) were higher in the TgF344-AD rats at 12 weeks. This also aligns with findings in human, from the AD Neuroimaging Initiative (ADNI), which observed DCA:CA ratios that were found to be significantly higher in the human AD group compared to normal older, early mild cognitive impairment, and also late mild cognitive impairment individuals [22].

Our study showed that certain lipid pathways seem to have been accelerated in TgF344-AD rats, leading to earlier and more notable metabolic changes than those observed in normal ageing. Previous studies have also highlighted other metabolic changes in AD, including disruptions in glucose metabolism in the brain— a hallmark of aging that occurs decades before clinical symptoms of AD appear. AD is associated with far more severe disruptions in glucose utilization, often described as “glucose hypometabolism” [36, 37].

Additionally, AD is linked to increased energy demands in neurons, which struggle to maintain function due to reduced glucose metabolism efficiency. This bioenergetic stress can lead to accelerated neuronal degeneration, in contrast to normal ageing, where neurons experience more modest metabolic changes [38]. Furthermore, AD also affects insulin signaling in the brain, with some researchers referring to it as “type III diabetes” due to its association with insulin resistance. This disrupted insulin signaling exacerbates metabolic stress, worsening the disease progression than the slower metabolic decline in normal ageing [39-41]. In summary, while normal ageing involves some metabolic decline, AD accelerates these processes, particularly in glucose metabolism, energy production, and insulin signaling, leading to a faster neurodegeneration and functional impairment. These changes, combined with lipid metabolic disruptions as highlighted in this study, suggest that AD disrupts multiple metabolic pathways simultaneously, creating a complex network of dysfunction that drives disease progression.

In this study, we identified 54 out of 825 lipids associated with sex, after adjusting for age. The ones most elevated in females were oleic acid, alpha-linoleic acid, 12-S-HEPE/12-HETE, PE(O-16:0/22:5), PC(16:1/18:0) and 15-HEPE/15-HETE. The ones most decreased in females were several TGs and PC(O-20:5/FA18:1). In males, specific bile acids or bile acid ratios (e.g. GCA, GUDCA, TDCA/DCA, GCA/GDCA) were downregulated in the AD group, while several SMs were upregulated in the AD group. These findings highlight sex-dependent lipid changes in the TgF344-AD rat model, providing novel insights into the metabolic differences between males and females and furthering our understanding of sex-specific AD pathology. Interestingly, our results show similar trends to those observed in a recent study from our laboratory using human ADNI samples [42]. In females, omega-3 derivatives (EPA and DHA) and omega-6 derivatives (arachidonic acid, AA) were found to be higher in AD compared to CN. Consistently, in our TgF344-AD rat model, 12-S-HEPE/12-HETE and 15-HEPE/15-HETE were elevated in females. Notably, 12-S-HEPE and 15-HEPE are derivatives of EPA, while 12-HETE and 15-HETE are derivatives of AA. These parallels between human and rat findings suggest that the TgF344-AD rat model serves as a robust translational model. This allows for the investigation of tissue-specific pathophysiology, testing interventions, or challenging models with inhibitors to zoom into disease processes and validate hypotheses on key steps involved in AD pathology. Furthermore, the model provides a platform for testing novel therapeutic strategies. Numerous studies in human have shown that women are disproportionately affected by AD, with both a higher incidence and faster cognitive decline compared to men [43]. It is thought that hormonal factors, particularly estrogen that declines in women after menopause, contribute to this increased susceptibility. Specifically, estrogen, an ovarian steroid hormone, is believed to influence lipid metabolism at various stages of biosynthesis, including playing a crucial role in lipid transport and exchange, enhancing the expression of metabolic enzymes, and reducing the oxidation of α-linoleic acid (α-LA) [44]. In AD rodent models, females have been shown to exhibit more severe cognitive deficits and higher levels of amyloid beta plaque accumulation compared to males [45]. Additionally, sex differences in brain metabolism [46], immune responses [47], and microglial activation [48, 49] have been observed in rodent studies, with females generally showing heightened neuroinflammation and oxidative stress [50]. Female rodents also tend to exhibit higher burden of amyloid plaques, neurofibrillary tangles, and tau pathology, all key markers of AD [51]. Lipids such as oleic acid, which plays a role in inflammatory processes, may contribute to these sex differences in neuroinflammation [52, 53]. Additionally, 12-S-HEPE/12-HETE and 15-HEPE/15-HETE, part of the eicosanoid pathway, have been reported to exhibit sex-specific differences due to the hormones influence [54, 55]. Other studies indicate that bile acid metabolism is sex-dependent, with women exhibiting different bile acid patterns compared to men. Estrogen is known to regulate bile acid synthesis and circulation, which could explain the sex-specific effects observed in AD patients [56-58]. Similarly, altered phospholipid metabolism, particularly in post-menopausal women, may affect the brain’s lipid composition and contribute to sex differences in AD [59]. Overall, while direct evidence on these specific metabolites in the context of sex differences in AD is still emerging, our findings underscore the influence of sex hormones and metabolism in modulating AD pathology. Further targeted studies are needed to better clarify these mechanisms. It should be noted that these are the first findings on sex related changes in lipid expression in the TgF344-AD rat plasma, providing novel insights into the model, as well as furthering our understanding of sex related AD pathology.

As our multilevel study included the variables age, sex and AD, it is challenging to evaluate each factor individually. Therefore, we developed multivariate models to discriminate between TgF344-AD and WT rats. Using stepwise regression, we developed the final predictive full age range and midlife models to assess the contribution of various covariates. Notably, these rat predictive models align with findings observed in human studies. In the full age range model, the most significant factor is age, followed by sex. This corresponds well with established knowledge that increasing age is the greatest risk factor for developing AD [60, 61]. Moreover, numerous studies have demonstrated the influence of sex, as females are disproportionately affected by AD, with both a higher incidence and faster cognitive decline compared to males [43]. Our full age range model also includes DHA, which has been observed to be decreased in the brain, blood, and CSF of AD individuals [59, 62-64]. Additionally, both 15-HETE and anandamide (AEA), included in the full age range model, highlight critical aspects of AD pathology. 15-HETE, linked to oxidative stress and inflammation being key factors in AD pathogenesis, has been found at elevated levels in the frontal and temporal cortices of AD patients, while lower levels of AEA have been observed in these same regions [65, 66]. We also developed a predictive midlife model for the 25-week and 50-week time points, but it performed less effectively than the full age-range model, which combines age, sex, and lipid variables. This reduced performance is likely due to the smaller sample size available when focusing only on these two time points, as splitting the data effectively halves the number of samples used for model training. This smaller dataset likely limits the model’s ability to capture the full pattern of lipid changes, resulting in lower accuracy and reliability compared to the combined model that includes the full age range. Interestingly, several lipids were found to overlap between the full age range model and the midlife model, specifically TDCA/DCA, 15-HETrE/DGLA, and LPG(16:1). This overlap suggests that these lipids may play a critical role across different stages of AD, making them potentially valuable targets for early intervention as well as for monitoring disease progression across the lifespan.

This study has several limitations. First, this study focused on lipidomic changes in the plasma of TgF344-AD and WT rats, while it does not consider other diseases. Therefore, it is possible that the observed changes in lipid profiles and other metabolites are not specific to AD. This raises the question of whether the lipidomic changes might occur as well in other neurodegenerative or metabolic processes. However, it is important to note that the plasma lipidomic fingerprint, which integrates a combination of both changed and unchanged lipids, could still provide a highly specific profile for AD, distinguishing it from other conditions. Another limitation is that, after adjusting the p-values using the Benjamini-Hochberg method, no statistically significant results were observed in the univariate analysis. However, clear trends were evident, and our multivariate analysis revealed biologically interesting patterns within the models. Additionally, this study is based on plasma samples and does not include data from the brain or CSF, which are critical in understanding AD pathology. However, from our longitudinal study, not only plasma but also CSF, brain tissue, and other tissues have been collected for future analysis. Analysis of these additional matrices will provide more comprehensive insights into how systemic changes in the plasma correlate with changes in the brain, CSF, and other body tissues. Additionally, the expression of transporters, which regulate lipid transport mechanism between the blood and brain, could be of interest to future research of better understanding the mechanisms involved in these metabolic changes. From the rats used in this study, also altered protein expression of membrane transporters in isolated cerebral microvessels and brain cortex have been identified [67], and these results can be integrated in future data analysis. A third limitation is that this is an animal study, and rats, while useful models, are not humans. There are inherent biological differences between species that must be considered when translating these findings to human AD pathology. Nevertheless, there are many interesting similarities between AD rat models and AD patients, as also earlier highlighted in Qin’s review [25]. These similarities allow animal models to remain valuable for studying AD mechanisms and potential treatments. One key advantage of using rat models is that they allow for longitudinal studies without the long timeframe required in human studies, while also allowing brain and other tissues to be collected. This is particularly beneficial for understanding the progression of AD over time and identifying potential therapeutic targets that may slow or halt disease progression. Thus, while this study has its limitations, it lays a solid foundation for future research, particularly when integrated with additional brain and CSF data. It serves as an essential step toward understanding and addressing AD progression.

## 5. Conclusion

In this rat study, comparing TgF344-AD and WT of male and female rats among different ages, we identified specific lipid alterations as potential early indicators of AD, paving the way for the development and evaluation of novel diagnostic tools and therapeutic targets. The full age-range and midlife predictive models provided fundamental insights into lipid alterations across life stages in both male and female TgF344-AD rats, highlighting distinct lipid signatures associated with AD. Moreover, the observed similarities in lipidomic profile changes between TgF344-AD rats and humans suggest potentially shared mechanistic pathways, offering a translational bridge between preclinical and clinical research. This finding highlights the value of lipidomics as a powerful tool not only for understanding AD onset and progression but also for evaluating the underlying mechanisms identified in human studies. Our study pioneers the application of lipidomics to the TgF344-AD rat model, offering a comprehensive framework for investigating lipid-based mechanisms in AD. These results provide a valuable foundation for better understanding of the interaction between lipid metabolism, aging, and neurodegeneration, as well as for assessing potential treatment strategies to address AD progression.

## Abbreviations list

AD: Alzheimer’s disease
TgF344-AD: transgenic rat model for AD
MCI: mild cognitive impairment
FDA: Food and Drug Administration
CSF: cerebrospinal fluid
APP: amyloid precursor protein
Aβ42: amyloid-β42
P-tau181: phospho-tau181
NfL: neurofilament light
PS1ΔE9: PSEN1 with a deletion of exon 9
WT: wildtype
MRM: multiple-reaction-monitoring
LLE: liquid liquid extraction
MTBE: methyl tert-butyl ether
AIC: Akaike Information Criterion
PCA: principal component analysis
PC1: the first principal component
PC2: the second principal component
TGs: triglycerides
PCs: phosphatidylcholines
PEs: phosphatidylethanolamines
SMs: sphingomyelins
GLM: generalized logistic model
CA: cholic acid
BAs: bile acids
LPAs: lysophosphatidic acids
ADNI: AD Neuroimaging Initiative
CN: normal older individuals
TCA: taurocholic acid
DCA: deoxycholic acid
TDCA: taurodeoxycholic acid
TCDCA: taurochenodeoxycholic acid
GCA: glycocholic acid
GDCA: glycodeoxycholic acid
GCDCA: glycochenodeoxycholic acid
GUDCA: glycoursodeoxycholic acid
EPA: eicosapentaenoic acid
AA: arachidonic acid.

## Acknowledgments

Chunyuan Yin acknowledges support from the China Scholarship Council fellowship (No. 202006550003). We thank the Predictive Pharmacology group for financing the animal work and providing the samples. We thank Biomedical Metabolomics Facility Leiden (BMFL) for their assistance with sample preparation and sample measurement.

## CRediT authorship contribution statement

**Chunyuan Yin**: Visualization, Writing – original draft, Writing – review and editing. **Alida Kindt**: Daily Supervision, Statistical methodology, Writing – review and editing. **Amy Harms**: Analytical methodology, Writing – review and editing. **Robin Hartman**: Animal experiments. **Thomas Hankemeier**: Supervision, Writing – review and editing. **Elizabeth de Lange**: Overall animal experimental design, overall supervision, Writing – review and editing.

